# Polyphenol oxidase silencing avoids protein crosslinking and enzymatic browning in *Nicotiana benthamiana* leaf extracts

**DOI:** 10.1101/2025.03.31.646341

**Authors:** Chidambareswaren Mahadevan, Emma C. Watts, Kaijie Zheng, Shi-jian Song, Renier A. L. van der Hoorn

**Affiliations:** The Plant Chemetics Laboratory, Department of Biology, University of Oxford, UK

**Keywords:** Polyphenol oxidase, browning, phenolics, agroinfiltration, *Nicotiana benthamiana*

## Abstract

Oxidation in extracts from agroinfiltrated leaves leads to browning and protein precipitation. Here we show that silencing of polyphenol oxidase (PPO) in *Nicotiana benthamiana* suppresses enzymatic browning and avoids crosslinking of endogenous proteins such as RuBisCo. *PPO* silencing does not impact plant growth or development and may also enhance transient gene expression. These results show that depletion of PPO activity could be highly beneficial for plant science applications and molecular pharming.

---

Browning of extracts during purification of recombinant proteins from agroinfiltrated leaves is a widely observed phenomenon that is commonly quelled with reducing agents and absorbing materials. In fruits and vegetables, browning results from the oxidation of phenolics into brown quinones which react with themselves and other molecules to form a brown melanin-like polymer (**Figure 1A**, Siu et al., 2023). The oxidation of phenolics is catalysed by polyphenol oxidase (PPO), which is localised to chloroplasts but oxidises vacuolar phenolics upon cell disruption. Enzymatic browning in sliced apple and bruised potato has been suppressed by silencing polypenol oxidase (PPO; Carter, 2013; Gonzalez et al., 2020).

**Figure 1.**
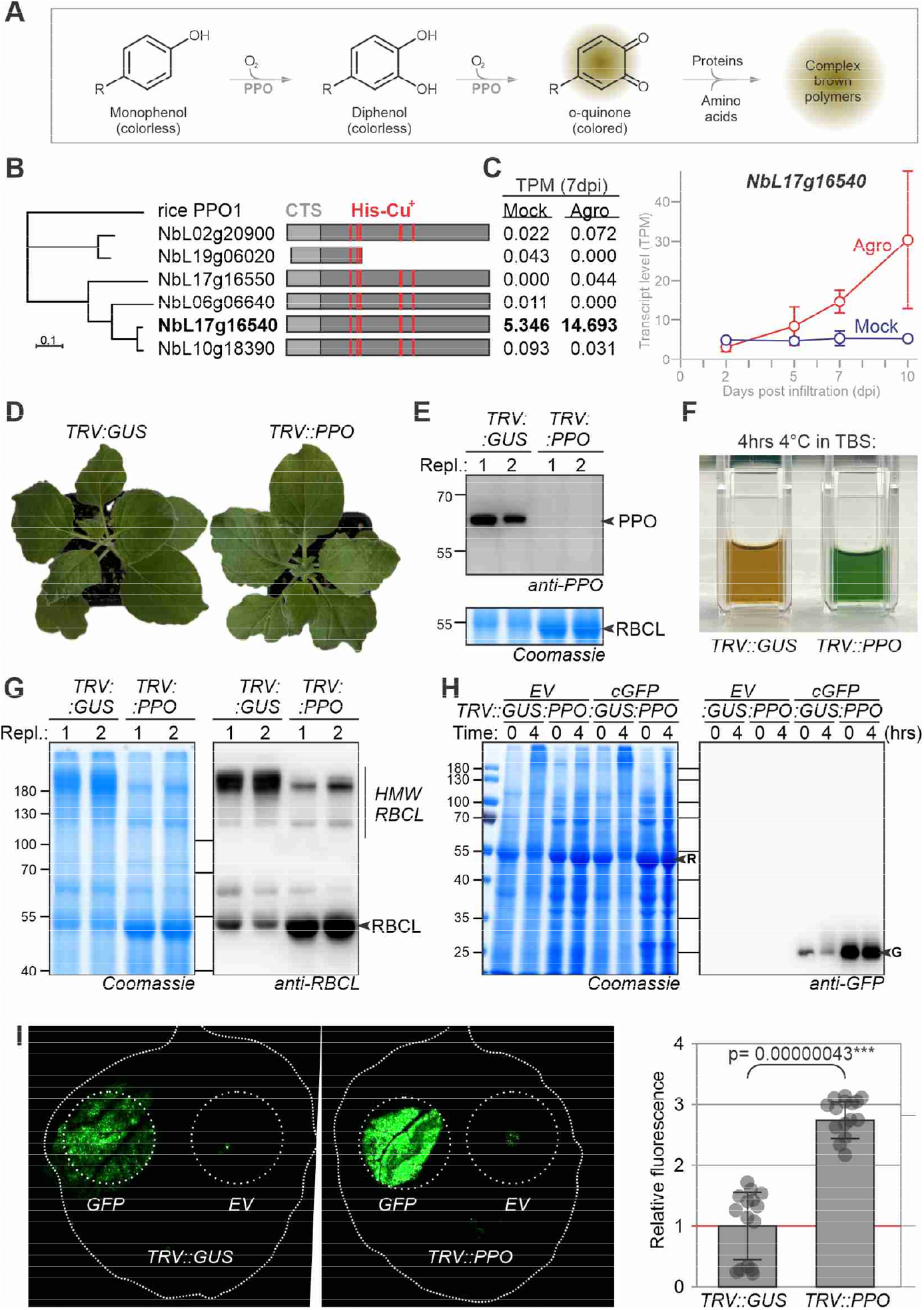
*PPO* silencing avoids browning and crosslinking and may increase transient expression. **A**) PPO oxidizes phenols into brown quinones, which react with themselves and with proteins, resulting in a brown polymer. **B**) Neighbor-joining phylogeny of six *N. benthamiana* PPOs, including rice PPO1 as the outgroup. Domain structures indicate chloroplast targeting signals (CTS) and catalytic His residues (red) that coordinate two copper ions that suspend a molecular oxygen. **C**) *PPO* transcript levels in transcripts per million reads (TPM) of mock- and agro-infiltrated leaves (from Grosse-Holz et al., 2018). Error bars represent SE of n=3 replicates. **D**) *PPO* silencing does not affect growth or development. 2-week-old plants were agroinoculated with tobacco rattle virus (TRV) carrying a 300bp fragment of GUS or PPO and images were taken 3 weeks later. **E**) PPO protein is depleted in *TRV::PPO* plants. Shown are two replicates. **F**) Total leaf extracts of *TRV::PPO* plants do not turn brown when incubated for 4hrs at 4°C in Tris-buffered saline (TBS). **G**) Crosslinking is reduced in leaf extracts of *TRV::PPO* plants when incubated for 4hrs at 4°C. (E-G) Total extracts of two replicates of 5-week-old *TRV::GUS* and *TRV::PPO* plants were incubated for 4hrs at 4°C, imaged in a cuvette (F), separated on protein gel and stained with Coomassie and analysed on western blot with anti-PPO antibody (E) or anti-RBCL antibody (G). R, RBCL; G, GFP. **H**) More GFP accumulates upon transient GFP expression in *TRV::PPO* plants. *TRV::PPO* and *TRV::GUS* plants were agroinfiltrated with pEAQ-HT-GFP-P19 and extracts were generated in TBS at 5dpi and incubated for 4hrs at 4°C and analysed by Coomassie staining and western blot with anti-GFP antibody. **I**) Increased GFP fluorescence in *TRV::PPO* plants. *TRV::PPO* and *TRV::GUS* plants were agroinfiltrated with 35S::GFP and imaged at 5dpi and fluorescence was quantified. Error bars represent SE of n=3 replicates. The p-value was determined with the Student’s t-test.

To prevent browning of extracts from agroinfiltrated leaves, we depleted *PPO* transcripts by virus-induced gene silencing (VIGS). The *N. benthamiana* genome (Ranawaka et al., 2023) contains six *PPO* genes, of which five encode putative functional enzymes (**Figure 1B**). However, only NbL17g16540 is significantly expressed in leaves and its expression increased upon agroinfiltration (**Figure 1C**). We cloned a 300bp fragment to silence NbL17g16540 and its homeolog (NbL10g18390, **Figure S1**) into RNA2 of the bipartite genome of Tobacco Rattle Virus (TRV). Young *N. benthamiana* plants were co-agroinoculated with TRV1 and TRV2 carrying fragments of *PPO* or GUS (β-glucuronidase, negative control). Three weeks post-infiltration, *TRV::PPO* plants showed no growth or developmental phenotypes compared to *TRV::GUS* plants (**Figure 1D**). Western blot analysis of leaf extracts from these plants confirms that PPO was successfully depleted from *TRV::PPO* plants (**Figure 1E**).

Importantly, cleared leaf extracts of *TRV::PPO* plants in Tris-buffered saline (TBS) remained green after 4 hours of incubation at 4ºC, in contrast to browning observed in extracts of *TRV::GUS* plants (**Figure 1F**). When these incubated extracts were separated on protein gels, *TRV::PPO* samples revealed much stronger signals at 55 kDa, whereas *TRV::GUS* samples showed high molecular weight (HMW) signals at >180 kDa (**Figure 1G**). Western-blot analysis revealed that much of the large subunit of Ribulose bisphosphate carboxylase large chain (RBCL) runs at HMW in the *TRV::GUS* sample, unlike the 55 kDa signal found in *TRV::PPO* samples (**Figure 1G**). This indicates that PPO catalyzes crosslinking of RBCL, possibly fixing RBCL tetramers in the multimeric RBCL complex (Duff et al., 2000). The HMW signal was less prominent at the t=0 time point (**Figure 1H**), indicating that crosslinking occurs during incubation.

To examine the impact of PPO depletion on protein expression and accumulation, we transiently expressed cytoplasmic GFP in *TRV::GUS* and *TRV::PPO* plants. Interestingly, extracts from *TRV::PPO* plants contained significantly higher levels of GFP protein than the *TRV::GUS* control plants (**Figure 1H**). Further, the intensity of the GFP signal is reduced in *TRV::GUS* extracts upon incubation for 4hrs at 4ºC but not in *TRV::PPO* plants. However, we did not detect HMW complexes containing GFP in *TRV::GUS* control plants (**Figure 1H**). Measuring GFP fluorescence directly from leaves revealed a significant, 2.8-fold higher GFP fluorescence from *TRV::PPO* plants when compared to *TRV::GUS* plants (**Figure 1I**), consistent with Western blot analysis and supporting the increased GFP accumulation in *TRV::PPO* plants. Although the underlying mechanism behind the increased transient expression is unclear, this feature might be a valuable advantage of *PPO* silencing.

Thus, PPO silencing avoids enzymatic browning and protein crosslinking and this may increase yield and quality of purified proteins, and improve routinely performed experiments such as co-immunoprecipitation and metabolic experiments. While *PPO* silencing does not affect plant growth or development, it may reduce immunity to pests and pathogens (Zhang & Sun, 2021). Alternative ways to deplete PPO activity include genome editing, the use of chemical or protein-based inhibitors, or physical methods such as used in the food industry (Sui et al., 2023).

## Acknowledgements

We would like to thank Ursula Pyzio for excellent plant care; Felix Homma for supporting RNAseq analysis; Sarah Rodgers, Caroline O’Brian and Patricia Bowman for technical support and George Lomonossoff for providing pEAQ vectors.

## Author contributions

RH conceived the project; CM performed most experiments with help of EW, KZ and SS; RH wrote the manuscript with help of all authors.

## Supplemental Materials

**Polyphenol oxidase silencing avoids protein crosslinking and browning in *Nicotiana benthamiana* leaf extracts** Chidambareswaren Mahadevan et al.

### Bioinformatic analysis

Transcriptomic data was processed using the High Performance Computing (HPC) compute clusters from the University of Oxford Advanced Research Computing facility (Richards, 2015). Raw sequence reads from a previous transcriptomics study (Grosse-Holz et al., 2018), were downloaded from Sequence Reads Archives (NCBI) using fasterq from the SRA toolkit v3.0.10 of NCBI. Quality trimming was performed with Trimmomatic v.039 (Bolger et al., 2014) and checked with FastQC v0.12.0. (Wingett and Andrews, 2018). Processed reads were then mapped to the LAB 3.6 genome (Ranawaka et al., 2023) using kallisto v0.50.1 (Bray et al., 2016). The phylogenetic tree of PPO genes was constructed with Geneious Prime® 2024.0.7 (https://www.geneious.com). A distance matrix was built with global alignment and free end gaps, employing the Blosum62 cost matrix. The tree was built using the Neighbor-Joining method with the Jukes-Cantor genetic distance model. *Oryza sativa* polyphenol oxidase I (Gene ID: 4337055) from NCBI was included as the outgroup.

### Plant cultivation conditions

Wild-type *Nicotiana benthamiana* (LAB) seeds were sown in a 3:1 mix of soil (Sinclair Modular Seed Peat reduced propagation mix) and vermiculite (Sinclair brand Pro Medium) in 7×7 cm square pots. Seeds were initially grown under high humidity, covered with transparent plastic for 5 days. After uncovering, seedlings were cultivated in the greenhouse at 80–120 µmol/m^2^/s light, with temperatures set to 21°C at night and 22–23°C during the day, under a 16-hour light cycle. Two-week-old plants were then agroinfiltrated with the respective TRV vectors and transferred to a controlled growth chamber set at 100 µmol/m^2^/s light, 21°C, and 50–60% relative humidity, also with a 16-hour light regime. Plants were watered three times per week, ensuring even moisture in the pots.

### Construction of plasmids

All plasmids used in this study are summarized in Supplemental **Table S1**. A 300 bp fragment of the *PPO* gene (Supplemental **Table S2**) was selected using the VIGS tool (Fernandez-Pozo et al., 2015), synthesized by Twist Biosciences, and cloned into the golden-gate compatible vector TRV2gg (Duggan et al., 2016) using a BsaI restriction enzyme reaction to generate expression plasmids. The resulting plasmid was transformed into *Escherichia coli* DH10β for amplification, purified, and subsequently transformed into *Agrobacterium tumefaciens* GV3101-pMP90. Transformants were selected on LB agar plates containing 25 µM rifampicin, 10 µM gentamycin, and 50 µM kanamycin. A single colony was cultured in liquid LB medium supplemented with the same antibiotics.

### Virus-induced Gene Silencing (VIGS)

*Agrobacterium* strains containing TRV1 or TRV2 (Supplemental **Table S2**) were grown overnight at 28°C in LB medium supplemented with 25 mg/L rifampicin, 10 mg/L gentamycin, and 50 mg/L kanamycin. Cultures were centrifuged at 3,500 x g for 10 minutes at room temperature, and the resulting pellets were resuspended in agroinfiltration buffer (10 mM MES, pH 5.7, 10 mM MgCl2, 100 µM acetosyringone) to an OD600 of 1.0. TRV1 and TRV2 cultures were then mixed in a 1:1 ratio. Two-week-old *Nicotiana benthamiana* plants were agroinfiltrated with this bacterial suspension using a 1 mL needleless syringe. Three to five weeks post-infiltration, plants were assessed for silencing by observing bleached leaves in *TRV::PDS* positive control plants. Successfully silenced plants were subsequently tested by further agroinfiltration.

### Protein extraction and incubation

Proteins were extracted from 1 cm leaf discs, which were punched from the same leaves as above and transferred to 1.5 mL Eppendorf tubes. Each tube received two steel balls (2.3 mm diameter, BioSpec) and 150 µL of extraction buffer (50 mM Tris-HCl, pH 7.6, 250 mM NaCl, 1 mM EDTA, 0.002% Tween-20). Protein extraction was performed using a QIAGEN TissueLyser II with the following settings: 30 strokes, 30 seconds per stroke, repeated twice. The samples were then centrifuged at 13,000 x *g* for 10 minutes at 4°C. Proteins were subsequently incubated in a cold room maintained at 4°C for the 4 hr time-course experiment.

### Western blot analysis

The total soluble protein supernatant was mixed at a 1:3 ratio with the 4x gel loading buffer (200 mM Tris-HCl, pH 6.8, 400 mM DTT, 8% SDS, 0.4% bromophenol blue, 40% glycerol) and heated at 95°C for 5 minutes. Proteins were separated on a 12% w/v SDS-PAGE gel, transferred onto a polyvinylidene difluoride (PVDF) membrane using the Trans-Blot Turbo system (Bio-Rad, Hercules, CA), and blocked for 1 hour at room temperature in 5% w/v skimmed milk in phosphate-buffered saline (PBS) with 0.01% v/v Tween-20. The blots were then incubated with primary antibodies: anti-PPO (1:2,500, MyBioSource) or anti-RBCL (1:2,500, Agrisera) or anti-GFP-HRP (1:3,000, Invitrogen) and with the secondary antibody, anti-rabbit (1:5,000, Invitrogen), in 5% w/v skimmed milk in PBS with 0.01% v/v Tween-20. Chemiluminescent signals were detected using the SuperSignal™ West Femto Maximum Sensitivity Substrate (Thermo Fisher Scientific, Waltham, MA, USA).

### GFP Expression, Imaging, and Quantification

Agrobacterium strain GV3101(pMP90) carrying the pEAQ-HT-GFP vector (encoding GFP and P19 along with an empty vector, EV) was grown overnight at 28°C in LB medium containing 25 µM rifampicin, 10 µM gentamycin, and 50 µM kanamycin. Cultures were centrifuged at 3,500 x g for 10 minutes at 21°C and resuspended in infiltration buffer (10 mM MES, 10 mM MgCl2, 100 µM acetosyringone, pH 5.7) to an OD600 of 0.5. Expanded leaves of virus-induced gene silencing (VIGS) plants were agroinfiltrated using a needleless syringe. At 5 days post-infiltration (dpi), leaves were detached and scanned for GFP fluorescence on an Amersham Typhoon 5 Biomolecular Imager (GE Healthcare Life Sciences, Little Chalfont, UK) using the 488 nm laser with Cy2 settings. The fluorescence quantification was performed using ImageJ software (Schneider et al., 2012). Statistical significance was assessed using Student’s *t*-test.

**Supplemental Table S1.** Used plasmids.

| Plasmid | Description | Reference |
| --- | --- | --- |
| *TRV::GUS* | Binary silencing construct for GUS | Duggan et al., 2021 |
| *TRV::PDS* | Binary silencing construct for PDS | Liu et al., 2002 |
| pCM02 | Binary silencing construct for PPO | This work |
| pEAQ-HT-P19 | Binary empty vector construct with P19 | Sainsbury et al., 2009 |
| pEAQ-HT-GFP | Binary expression construct for GFP + P19 | Sainsbury et al., 2009 |

**Supplemental Table S2.**
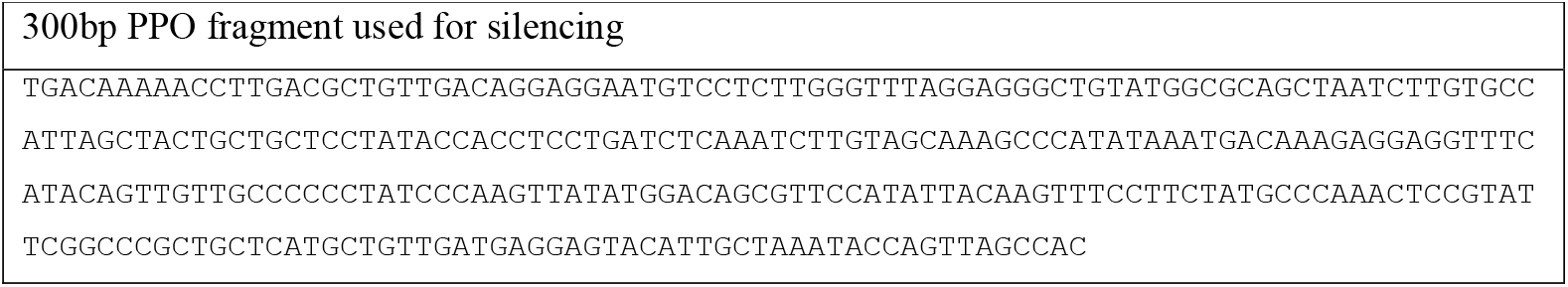
Oligonucleotides.

**Figure S1.**
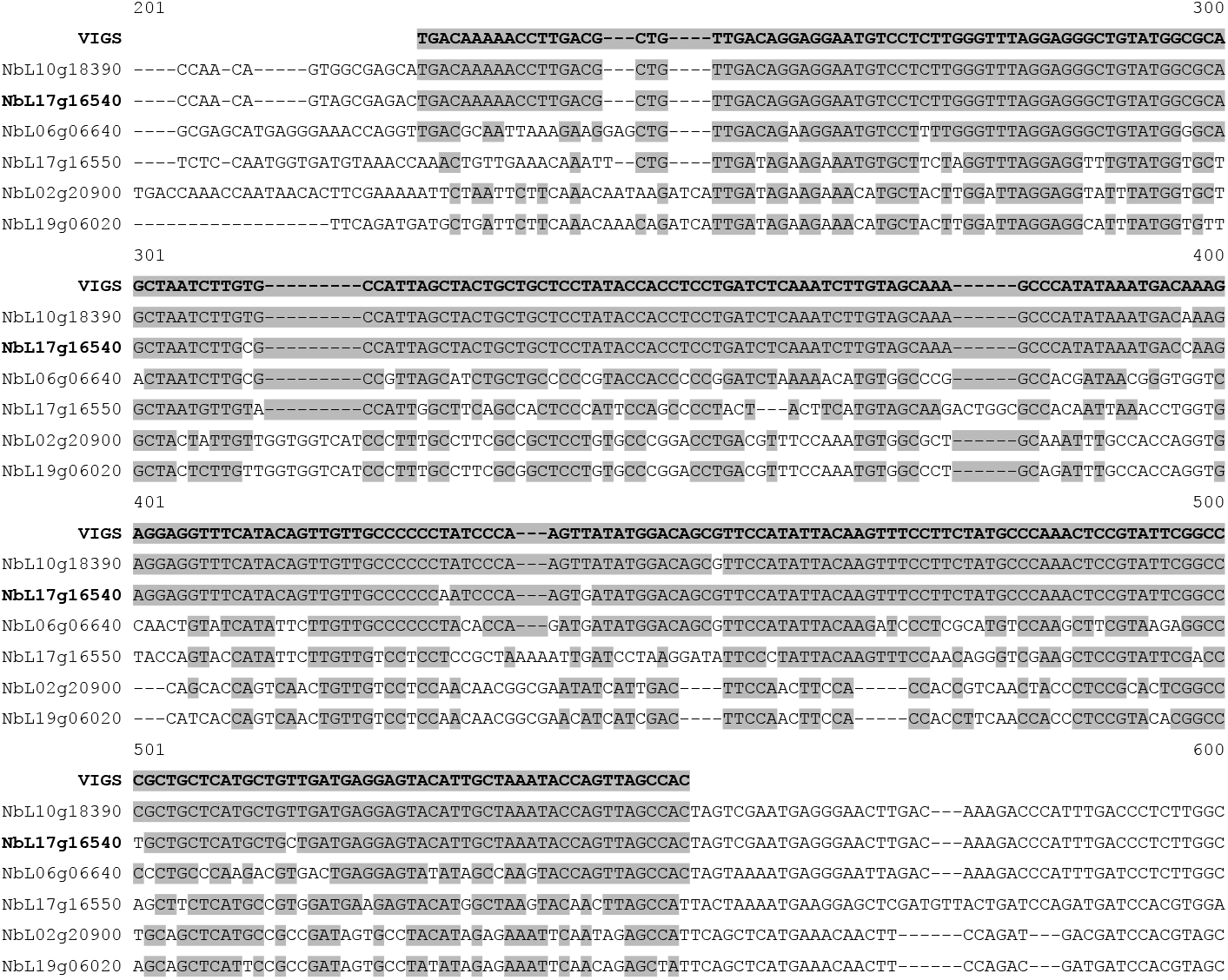
Alignment of VIGS fragment with 6 *PPO* genes of *N. benthamiana*

## References

Carter N. (2012) Petition for determination of nonregulated status: arctic™ apple (Malus x domestica) Events GD743 and GS784, United States Department of Agriculture—Animal and Plant Health Inspection Service, 2012, available from: https://www.aphis.usda.gov/brs/aphisdocs/10_16101p.pdf.

Dodds, I., Chen, C., Buscaill, P. and van der Hoorn, R.A.L. (2023) Depletion of the NbCORE receptor drastically improves agroinfiltration productivity in older Nicotiana benthamiana plants. Plant Biotechnol. J. 21, 1103–1105.

Duff AP, Andrews TJ, Curmi PM. (2000) The transition between the open and closed states of rubisco is triggered by the inter-phosphate distance of the bound bisphosphate. J. Mol. Biol. 298:903–916.

González MN, Massa GA, Andersson M, Turesson H, Olsson N, Fält AS, Storani L, Décima Oneto CA, Hofvander P, Feingold SE (2020) Reduced enzymatic browning in potato tubers by specific editing of a polyphenol oxidase gene via ribonucleoprotein complexes delivery of the CRISPR/Cas9 system. Front Plant Sci. 10:1–12.

Ranawaka B, An J, Lorenc MT, Jung H, Sulli M, Aprea G, Roden S, Llaca V, Hayashi S, Asadyar L, LeBlanc Z, Ahmed Z, Naim F, de Campos SB, Cooper T, de Felippes FF, Dong P, Zhong S, Garcia-Carpintero V, Orzaez D, Dudley KJ, Bombarely A, Bally J, Winefield C, Giuliano G, Waterhouse PM. (2023) A multi-omic Nicotiana benthamiana resource for fundamental research and biotechnology. Nat. Plants. 9:1558–1571.

Sui X, Meng Z, Dong T, Fan X, Wang Q. (2023) Enzymatic browning and polyphenol oxidase control strategies. Curr. Opin. Biotechnol. 81:102921.

Zhang J, Sun X. (2021) Recent advances in polyphenol oxidase-mediated plant stress responses. Phytochemistry. 181:112588.

## Supplemental References

Bolger, A. M., Lohse, M., & Usadel, B. (2014). Trimmomatic: A flexible trimmer for Illumina sequence data. Bioinformatics 30, 2114–2120.

Bray, N. L., Pimentel, H., Melsted, P., & Pachter, L. (2016). Near-optimal probabilistic RNA-seq quantification. Nat. Biotechn. 34, 525–527.

Dodds I, Chen C, Buscaill P, van der Hoorn RAL. (2023) Depletion of the NbCORE receptor drastically improves agroinfiltration productivity in older Nicotiana benthamiana plants. Plant Biotechnol. J. 21, 1103–1105.

Duggan C, Tumlas Y, Bozkurt TO. (2021) A golden-gate compatible TRV2 virus induced gene silencing (VIGS) vector. Zenodo 10.5281/zenodo.5666891.

Fernandez-Pozo N, Rosli HG, Martin GM, Mueller LA. (2015) The SGN VIGS tool: user-friendly software to design virus-induced gene silencing (VIGS) constructs for functional genomics. Mol. Plant 8, 486–488.

Grosse-Holz, F., Kelly, S., Blaskowski, S., Kaschani, F., Kaiser, M., & Van Der Hoorn, R. A. L. (2018). The transcriptome, extracellular proteome and active secretome of agroinfiltrated Nicotiana benthamiana uncover a large, diverse protease repertoire. Plant Biotechn. J. 16, 1068–1084.

Kourelis J, Marchal C, Posbeyikian A, Harant A, Kamoun S. (2023) NLR immune receptor-nanobody fusions confer plant disease resistance. Science 379, 934–939.

Liu Y, Schiff M, Dinesh-Kumar SP. (2002) Virus-induced gene silencing in tomato. Plant J. 31, 777–786.

Ranawaka, B., An, J., Lorenc, M. T., Jung, H., Sulli, M., Aprea, G., Roden, S., Llaca, V., Hayashi, S., Asadyar, L., LeBlanc, Z., Ahmed, Z., Naim, F., De Campos, S. B., Cooper, T., De Felippes, F. F., Dong, P., Zhong, S., Garcia-Carpintero, V., Orzaez, D., Dudley, K. J., Bombarely, A., Bally, J., Winefield, C., Giuliano, G., Waterhouse, P. M. (2023). A multi-omic Nicotiana benthamiana resource for fundamental research and biotechnology. Nat. Plants 9, 1558–1571.

Richards, A. (2015). University of Oxford Advanced Research Computing.

Sainsbury F, Thuenemann E C, Lomonossoff G P. (2009) pEAQ: versatile expression vectors for easy and quick transient expression of heterologous proteins in plants. Plant Biotechnol. J. 7, 682–693.

Schneider C A, Rasband W S, Eliceiri K W. (2012) NIH Image to ImageJ: 25 years of image analysis. Nat. Methods 9, 671–675.

Wingett, S. W., & Andrews, S. (2018). FastQ Screen: A tool for multi-genome mapping and quality control. F1000Research 7, 1338.

